# Thermoregulatory Constraints on Regenerative Competence: Evolutionary Trade-Offs Between Metabolic Homeostasis and Tissue Repair

**DOI:** 10.64898/2026.03.09.710613

**Authors:** Daniel Pelaez, Chloe Moulin, Jaelyn Chang, Kendall W. Knechtel

**Affiliations:** Bascom Palmer Eye Institute, Department of Ophthalmology, University of Miami Miller School of Medicine, Miami, FL, 33136, USA; Department of Cell and Systems Biology, University of Miami Miller School of Medicine, Miami, FL, 33136, USA; University of Miami Miller School of Medicine, Miami, FL, 33136, USA

**Keywords:** Thermoregulation, Regeneration, Endothermy, Central Nervous System, Evolution, Calcium, Endo/Sarcoplasmic Reticulum, SERCA

## Abstract

Regenerative capacity varies widely across the animal kingdom, yet adult mammals and birds exhibit limited ability to regenerate complex tissues such as the central nervous system (CNS). The evolutionary basis for this loss of regenerative competence remains poorly understood. Here we advance a Thermoregulatory Theory of Regenerative Scope, proposing that the emergence and refinement of endothermic physiology shaped tissue-specific molecular architectures that can bias injury responses away from regeneration in highly metabolic organs. Rather than imposing a systemic energetic constraint, we suggest that endothermy provided the evolutionary context in which thermogenic calcium-handling systems became increasingly specialized in excitable tissues. In particular, futile Ca²⁺ cycling across the sarco/endoplasmic reticulum through SERCA pumps contributes to heat generation but may also promote sustained intracellular Ca²⁺ elevations that activate inflammatory and fibrotic/gliotic signaling pathways incompatible with functional regeneration. Comparative observations indicate that regenerative competence is retained in ectothermic vertebrates and in mammalian contexts where thermoregulatory systems are developmentally immature or physiologically attenuated. Importantly, regenerative outcomes differ substantially among tissues within endothermic organisms, suggesting that local calcium regulatory architectures, rather than metabolic intensity alone, may determine regenerative potential. This framework generates experimentally testable predictions linking thermogenic Ca²⁺ signaling to regenerative failure and provides an evolutionary lens through which disparate observations in regenerative biology may be unified.

## Introduction

Within the breadth and scope of more than 4.5 billion years of evolutionary refinement by natural selection, the molecular mechanisms underlying most biological processes can be found and studied. This extensive evolutionary molecular repertoire offers a unique opportunity to elucidate potential solutions to intractable challenges in human health, such as the negligible regenerative capacity of the central nervous system (CNS). Regeneration – conceptualized as the ideal outcome of a healing process – varies widely across species throughout the animal kingdom, ranging from what is categorized as biological ‘immortality’ in some non-vertebrates (e.g. *Turritopsis dorhnii* and *Hydra vulgaris*) [1, 2], to full regenerative competence of limbs and CNS structures in fish and amphibians[3–6]. Unlike other vertebrates, however, most species within the mammaliaform and avian clades undergo regeneration failure following injury to the CNS [7, 8]

The questions of why and how some vertebrate species retain robust regenerative competence while others, particularly mammals and birds, lost this capacity have eluded evolutionary biologists and biomedical researchers for decades [9, 10]. Answers to these questions can unlock critical insights for regenerative medicine and human health.

Despite the general inability to regenerate, there exist instances within the mammalian and avian clades in which robust CNS regeneration can still be observed. Immediately after its development, and approximately up to the first postnatal week in rats and mice, as well as in the chicken while *in ovo*, the optic nerve can undergo spontaneous regeneration when injured [11–14]. The naked mole rat (*Heterocephalus glaber*), while possessing a regressed visual system that senses light presumably only for circadian and seasonal rhythm entrainment [15], undergoes spontaneous regeneration of the axons in its optic nerve following crush injury[16]. The naked mole rat has long been a model organism for mammalian aging and cancer research due to its unusually long lifespan for a rodent, negligible cellular senescence[17, 18], and extremely low incidence of neoplastic disease[19]. Recently, other rodent species – the brush-furred rat (*Lophuromys*), and the spiny mouse (*Acomys*) – have garnered attention as regenerative models for mammals [20, 21], including for the spiny mouse’s increased capacity for retinal regeneration compared to the traditional laboratory mouse (*Mus musculus*) [22, 23]. Understanding what drives the enhanced regenerative potential in altricial mammals and birds, the naked mole rat and *Acomys*, and whether there is any mechanistic overlap with the robust CNS regenerative competence seen in fish and amphibians would significantly advance our approach to promoting enhanced healing outcomes in humans. These instances establish a framework in which elucidating conserved mechanisms governing regenerative competence can be performed through comparative biology across clades, where large differences exist across all domains of molecular biology, between species of the mammalian clade, where these differences are lessened, and within individuals of a particular species (Figure 1).

**Figure 1:**
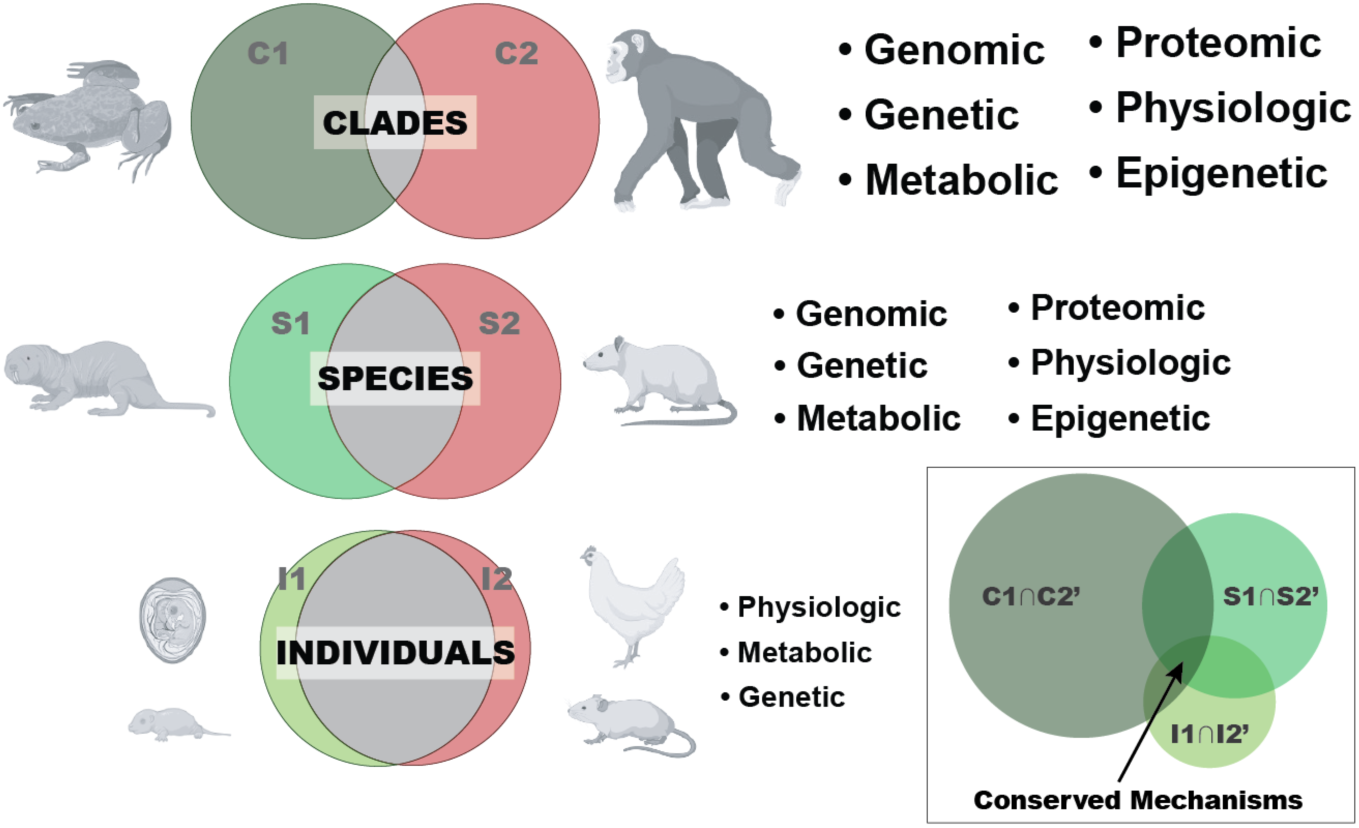
Schematic of the comparative biology framework, molecular biology domains in which differences are expected in each comparator group, and regenerative segment intersection in which conserved mechanisms underlying enhanced regenerative competence can be elucidated

## Thermoregulation and Regeneration

Upon closer analysis, a notable conserved mechanism that emerges between the regenerative segments in the above-highlighted examples relates to the maturity and complexity (or lack thereof) of the molecular mechanisms responsible for internal body heat generation (endothermy) and thermoregulation. While this is clear for fully ectothermic vertebrate species such as reptiles, amphibians, and fish, the link between whole-body endothermy and regenerative competence becomes much more nuanced in birds and mammals due to the myriad mechanisms underlying endothermic and homeothermic control, as well as the difference in developmental timing and maturity of these mechanisms between species (and even within individuals) of these clades (Figure 2A).

**Figure 2:**
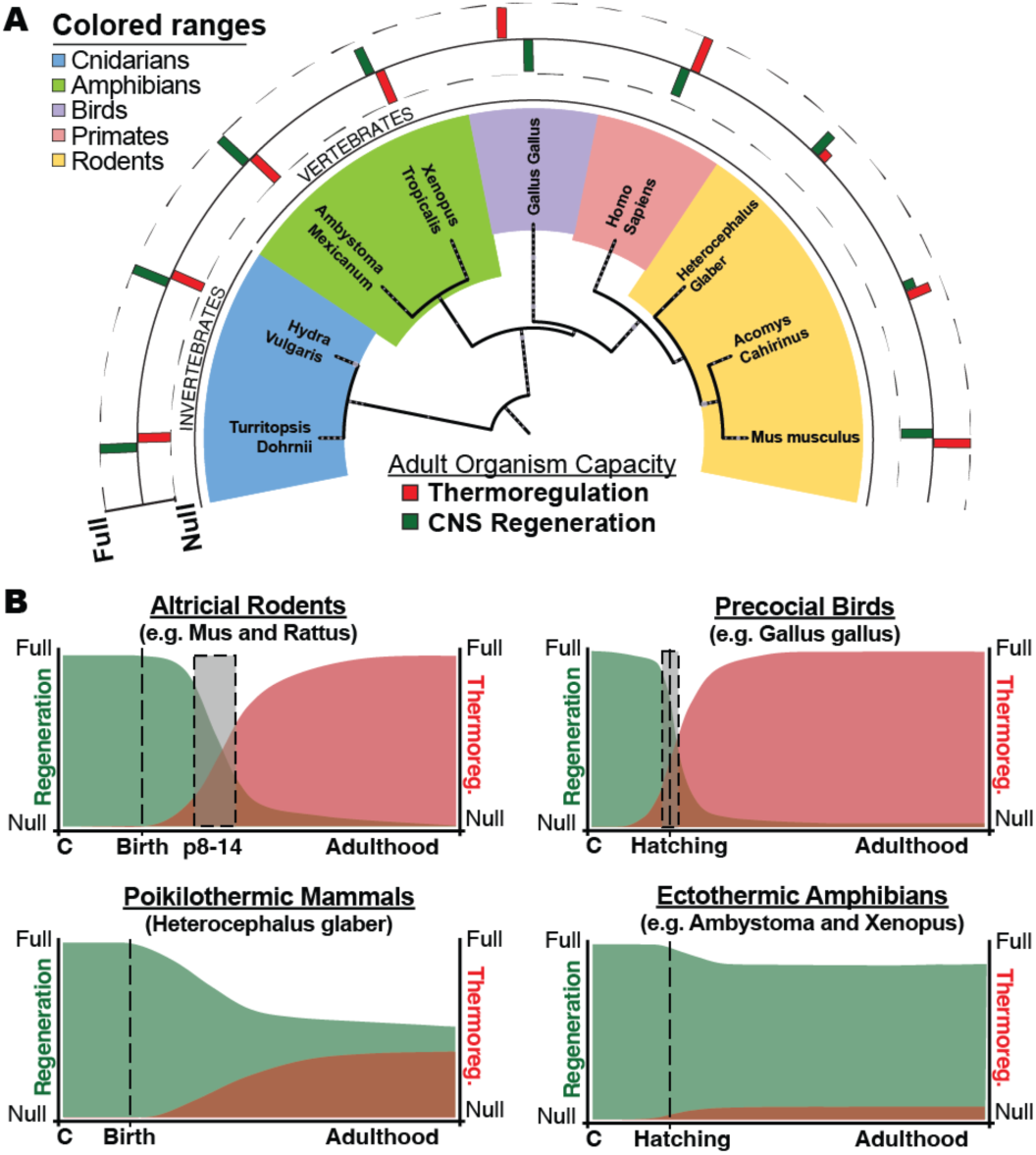
(A) Phylogenetic tree of species across the animal kingdom with disparate nervous regeneration and thermoregulatory capacities. (B) Schematic depicting the temporal counterbalance between the acquisition of thermoregulatory maturity and the regenerative capacity in different vertebrate groups.

Within the mammalian clade, only the naked mole rat is recognized as a true poikilotherm (i.e. an organism that cannot regulate its own body temperature except by behavioral means) with weak endothermic capacity, adding to the potential argument for a counterbalance between the molecular mechanisms for thermoregulation and regenerative competence. Moreover, while generally recognized as endothermic homeotherms, the thermoregulatory capacity of several members of the *Acomys* and *Lophuromys* families has been shown to be highly dependent on their environment [24–27] indicating a significant degree of poikilothermy, and inadequate or underdeveloped homeothermic mechanisms in these rodent species. A recent study in *Lophuromys melanonyx* [26] demonstrated a decreased thermoregulatory mechanism in the alpine species of these brush-furred rats resulting in a lower heat insulation capacity and suggested a need for solar radiation to support full homeothermy in these diurnal animals. In both *Acomys Russatus* as well as *Acomys Cahirinus*, significant variations in thermal regulation and metabolic rate adaptative capacity to varying environmental conditions have been recorded between populations of animals captured from different habitats [24, 27], showing a degree of poikilothermy which has not yet been studied in detail.

In mice and chicken, a central developmental timing difference in thermoregulatory capacity relates to the precocial and altricial state of these species at birth or hatching, respectively. Embryonically, both species are ectothermic and poikilothermic, relying on external heat sources (namely placental or *in ovo* incubation) without which the embryo is non-viable. As precocial birds, however, chicken transition to an endothermic phenotype upon hatching [28–30], whereas mice and most mammals are altricial at birth, and continue to require external thermal support for several days postnatally [31–33]. Laboratory mouse pups (*Mus musculus*) remain fully poikilothermic for the first week of life and do not develop endothermic and homeothermic regulation until around postnatal day 15 [34]. In rats, full physiological thermoregulatory abilities through both chemical and physical mechanisms is achieved by 14 to 18 days after birth [35]. The remarkable temporal overlap between the appearance and maturation of endothermic and thermoregulatory mechanisms and the decline in regenerative capacity within birds and mammals is highly conspicuous yet remains purely circumstantial (Figure 2B). While other researchers have highlighted this observation in other organ systems and offered mechanistic hypotheses [36], whether these are valid and a causative relationship exists remains to be proven.

## Thermoregulatory Theory of Regenerative Scope

Teleologically, the loss of regenerative capacity appears counterintuitive to survival, as regeneration enhances resilience to injury. Arguably however, the evolution of endothermy itself is equally as counterintuitive as one of the most energetically expensive biological processes in vertebrates, equivalent to injury response and healing processes, and surpassed only by growth during development [37, 38]. Several hypotheses explain the evolution of endothermy, with the aerobic scope hypothesis – positing that endothermy enhances locomotion and metabolic capacity – being the most widely accepted [38, 39]. A proposed byproduct of endothermy is the development of larger brains and increased neuronal density, which can confer cognitive advantages that offset the loss of regenerative competence [40]. Hypotheses linking thermoregulation to regenerative scope, however, remain largely observational and not testable.

To begin exploring these argumentatively, we consider a proof by contradiction: Could mammals and birds have evolved and thrived without endothermy? Ectothermic organisms, reliant on environmental heat, face increased predation risk during thermoregulation, necessitating robust healing mechanisms to survive non-lethal injurious encounters. In contrast, endothermy enables night foraging and hiding, habitation of resource-diverse environments, and reduced predation, potentially relieving evolutionary pressure to maintain full regenerative competence. This is further supported by the fact that endothermic species produce fewer offspring compared to ectotherms, as the endothermic survival adaptation can prioritize quality over quantity [41]. Consistent with this, the naked mole rat has one of the highest reproductive rates for a mammal and is recognized as a unique exception to the ‘one-half rule’ for their remarkably large litter sizes [42].

Under a more axiomatic proof, we accept endothermy to be fundamentally predicated on sustained high metabolic flux and make the assumption that an enhanced healing capacity (up to and including full tissue regeneration) confers improved somatic maintenance and thereby extends lifespan. *Ipso facto*, species with lower baseline metabolic rates and reduced thermogenic demands would be predicted to exhibit both slower aging trajectories and greater longevity. This hypothesis is consistent with empirical observations in long-lived mammals characterized by comparatively low metabolic intensity and low core temperatures. The longest-lived mammal, the bowhead whale, demonstrates exceptionally low mass-specific metabolic rates and an average deep body temperature lower than that of most non-hibernating eutherian mammals [43, 44]. Similarly, long-lived terrestrial mammals such as the two-toed sloth and the naked mole-rat exhibit reduced metabolic rates and significantly longer lifespans relative to phylogenetically comparable species. Collectively, these data support the working model that diminished thermoregulatory metabolic demand may facilitate extended lifespans through enhanced healing competences.

How, then, could thermoregulatory complexity interfere with regenerative competence? Building upon our conceptual framework, our theory delineates two interdependent mechanistic dimensions that can collectively explain this inverse functional relationship:

### Energetic resource constraints

Heat generation and maintenance in endothermic homeotherms is arguably the most metabolically demanding process in vertebrate biology, rivaled only by healing, growth, and regeneration. The high energetic cost of homeothermy may limit the availability of bioenergetic resources for functional healing (regeneration), particularly in the most metabolic demanding tissues of a ‘warm-blooded’ organism, such as the myocardium or CNS [45]. This competition for energy can explain why regenerative processes in the most metabolically demanding tissues are curtailed in mammals as their endothermic capacity expands [36]. Studies in laboratory mice have suggested that maturation of the central nervous system is probably the main factor responsible for the development of thermogenesis [34]. In the myocardium, the temporal correlation between the loss of regenerative capacity and the development of endothermy has been linked to changes in thyroid hormone levels around the same developmental period [36]. Notably, metabolic intensity alone cannot explain regenerative capacity across tissues. Several organs with high basal metabolic activity retain substantial regenerative competence, whereas others with comparable energetic demands do not. This observation suggests that regenerative limitations are unlikely to arise from global energetic constraints alone. Instead, the relevant determinants may lie in tissue-specific molecular frameworks that arose in the evolution of thermogenesis and conflict with injury response and healing processes such as cellular plasticity following injury.

### Molecular crosstalk interference

The molecular mechanisms underlying thermogenesis directly interfere with injury response and regenerative pathways in mammals. Heat is generated when any amount of free energy is not converted into downstream chemical or kinetic energy (work) but rather dissipated into the surroundings. Physiologically, thermogenesis results from the decoupling of the release of chemical energy (stored in ATP and other bioenergetic molecules) from the establishment of the chemiosmotic gradients (work) that drive myriad cell biological processes. This decoupling and loss of chemiosmotic control can disrupt the tightly orchestrated processes required for regeneration, including cell cycle reentry and transient proliferation, cellular reprogramming, and/or tissue remodeling [46–48].

## Thermoregulatory and Regenerative Mechanism Crosstalk

Endothermy is an umbrella term which encompasses a multitude of physical and biochemical mechanisms for internal heat generation in different species, and even within an individual. In broad mechanical terms, endothermic mechanisms are categorized into shivering thermogenesis and non-shivering thermogenesis (NST). Biochemically, however, endothermic mechanisms can be categorized into two different types: *i.* those that involve the flux of protons (H^+^), and *ii.* those that involve the flux of charged ions — most notably free calcium ions (Ca^2+^) — when uncoupled from the chemical energy expenditure intended to drive their compartmentalization in the cell. The most well-known example of NST involving proton movement occurs in the mitochondria of brown adipose tissues (BAT), where the permeation of the inner mitochondrial membrane (IMM) to protons (H^+^) by uncoupling proteins (UCP1/2/3) results in futile proton shuttling at the expense of NADH and FADH_2_ oxidation by complexes of the electron transport chain (ETC) and dissipation of the excess decoupled energy as heat. In shivering thermogenesis, rapid, small, involuntary muscle contractions in response to cold temperatures are triggered by calcium ion (Ca^2+^) release from sarcoplasmic reticulum (SR) stores and their subsequent binding to troponin to enable the establishment of the myosin-actin cross-bridge. Detachment of the myosin head from actin filaments requires ATP hydrolysis to prepare the contractile unit for the next interaction. Unlike in large coordinated and voluntary muscle contractions, during shivering thermogenesis, restriction of calcium reuptake to the SR through the Sarcoplasmic/Endoplasmic Reticulum Calcium ATPase (SERCA) by negative regulators like sarcolipin (SLN) [49] and phospholamban (PLN) [50] uncouples ATP hydrolysis from Ca^2+^ flux [51], resulting in multiple subsequent, and uncoordinated small muscle contractions and heat generation. Shivering thermogenesis thus results in significant heat generation through ATP hydrolysis at two sites: the myosin head uncoupled from force-generating muscle movements (work), and at SERCA channels uncoupled from calcium reuptake (work) to the SR/ER stores.

Calcium is a ubiquitous second messenger, and calcium ion fluxes underlie a vast number of biological processes, making the elucidation of molecular crosstalk interference between thermoregulation and regeneration a difficult task. Central to the discussion, however, sustained elevation of intracellular Ca²⁺ is a central proximal signal linking tissue injury to inflammatory amplification and fibrotic or gliotic remodeling. Cytosolic Ca²⁺ overload arising from plasma membrane influx or endoplasmic reticulum (ER) release through IP₃ (IP_3_R) and ryanodine receptors (RyR) activates multiple kinase pathways converging on transcriptional programs governed by NF-kappaB that drive expression of pro-inflammatory mediators [52–54]. In parallel, Ca²⁺ mobilization promotes assembly and activation of the NLRP3 inflammasome, facilitating caspase 1-dependent maturation of IL-1β and IL-18 and propagating sterile inflammation [55, 56]. Persistent Ca²⁺ signaling also potentiates profibrotic pathways by augmenting transforming growth factor beta (TGF-β)/SMAD signaling, reinforcing extracellular matrix deposition [55]. In the adult mammalian CNS, pervasive intracellular Ca²⁺ overload has been demonstrated as an injury-induced mediator of mitochondrial oxidation and structural remodeling [57]. Moreover, Ca^2+^ waves in astrocytes enhance inflammatory and STAT3-dependent transcriptional responses, promoting reactive gliosis characterized by GFAP upregulation and glial scar formation [58–60]. Conversely, in the regenerative competent *xenopus laevis* [61], Ca^2+^ transients following optic nerve transection are short-lived, and calcium dynamics return to homeostatic levels within minutes. Inhibition of SERCA2 — but not Na^2+^ channels — in this model extends calcium permanence and slows clearance and abrogates the regenerative outcome and functional recovery of sight compared to the SERCA2-unihibited controls [manuscript in preparation]. In a recent review, the strategies that promote axonal regeneration in the adult mammalian CNS have been shown to modulated intracellular calcium dynamics via one of three mechanisms: *i.* restriction of Ca^2+^ release from the ER via IP_3_R and RyR; *ii*. sequestration or chelation of free Ca^2+^ ions, or *iii*. Stimulation of Ca^2+^ reuptake to ER stores via modulation of SERCA negative regulators, with strategies that engage more than one of these mechanisms exerting a synergistic effect on axonal regeneration [manuscript in preparation]. Collectively, these data position dysregulated and sustained intracellular free Ca²⁺ as a nodal integrator of inflammatory transcriptional activation and maladaptive fibrotic or gliotic remodeling across organ systems in mammals.

Tissue-specific differences in calcium signaling architectures further support this mechanistic axis. Calcium dynamics are subject to stringent spatial and molecular regulation across tissues within an individual and between species, with distinct repertoires of sarco/endoplasmic reticulum Ca²⁺-ATPase (SERCA) modulators shaping both metabolic output and cellular fate decisions [62, 63]. The central nervous system (CNS), which by mass is the most metabolically active organ system in mammals and a major contributor to endothermic heat production [45], exemplifies this specialization. In excitable tissues, futile Ca²⁺ cycling across the endoplasmic reticulum (ER) constitutes a thermogenic mechanism that elevates basal metabolic rate. Notably, hepatocytes retain robust regenerative capacity despite belonging to one of the most metabolically active organs in mammals. Importantly, hepatocytes lack functional ryanodine receptors (RyR) and rely primarily on inositol trisphosphate receptor (IP₃R)-mediated Ca²⁺ release for intracellular calcium signaling [64, 65]. This architecture restricts the rapid and large-amplitude Ca²⁺ release events characteristic of RyR-dominated excitable tissues such as neurons, cardiomyocytes, and skeletal muscle. Consequently, hepatocytes may be less susceptible to sustained cytosolic Ca²⁺ elevations that promote inflammatory amplification and fibrotic remodeling following injury. From this perspective, the regenerative competence of the liver does not contradict the thermoregulatory framework. Rather, it supports the idea that specific calcium regulatory architectures, rather than metabolic intensity per se, may modulate regenerative outcomes. Skeletal muscle provides an important nuance to this framework. Striated muscle expresses ryanodine receptors and constitutes the primary site of shivering thermogenesis in mammals, processes that involve rapid Ca²⁺ release from sarcoplasmic reticulum stores. Despite this thermogenic architecture, skeletal muscle retains substantial regenerative capacity throughout adult life. This apparent exception can be reconciled by considering the cellular basis of muscle repair. Regeneration of skeletal muscle is mediated predominantly by satellite cells, a population of resident stem/progenitor cells capable of proliferating and differentiating into new myofibers following injury [66, 67]. In this context, regeneration does not require mature myofibers themselves to reverse terminal differentiation or reenter the cell cycle. Instead, repair occurs through progenitor-cell mediated replacement of damaged tissue. This distinction suggests that tissues whose regeneration depends on dedifferentiation or cell-cycle reentry of post-mitotic specialized cells (such as neurons or cardiomyocytes) may be particularly vulnerable to Ca²⁺-dependent inflammatory amplification and fibrotic signaling, whereas tissues capable of stem-cell mediated replacement may remain regenerative despite thermogenic specialization.

Unlike in striated muscle, canonical SERCA regulators such as phospholamban (PLN), myoregulin (MRLN), and sarcolipin (SLN) have not been identified in the mammalian CNS, implying the presence of distinct neural modulators. Neuronatin (NNAT), a neural tissue ER-associated proteolipid, has been linked to neurogenesis and systemic metabolic regulation [63, 68] positioning it as a candidate integrator of developmental and thermogenic programs in these tissues. A homology-based screen of SERCA-interacting proteins identified the transmembrane domain of the Tripartite Motif Containing 59 (TRIM59) protein as structurally and molecularly analogous to SLN (Figure 3A-C), raising the possibility that TRIM59 modulates ER Ca²⁺ handling in neural tissue. Intriguingly, NNAT lacks clear orthologs in regeneration-competent species, and the transmembrane domain of TRIM59 is significantly divergent in these organisms (Figure 3D). Finally, protein docking modeling shows a strong interaction between human SERCA2 and the transmembrane-domain of the human TRIM59 (Figure 3E), whereas there were no predicted interactions between these proteins from *xenopus laevis* (not shown).

**Figure 3:**
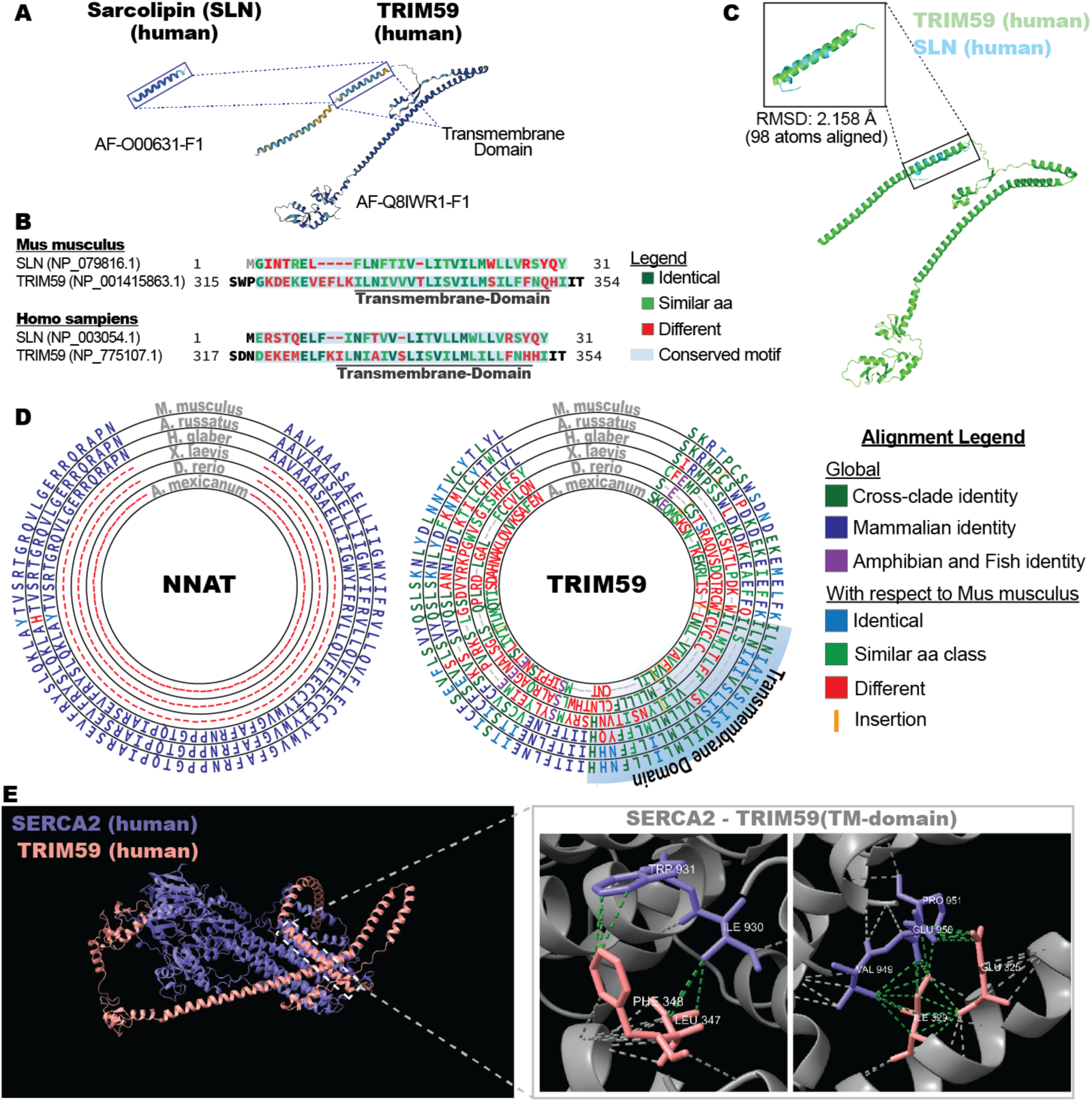
Structural (A) and amino acid sequence (B) homology between sarcolipin (SLN) and the transmembrane-domain (TM) of the tripartite motif containing 59 (TRIM59) protein in humans and mice. (C) Atomic positional alignment between the TM-TRIM59 and SLN from humans. (D) Sequence alignments of neuronatin (NNAT) and TRIM59 across species with divergent regenerative capacity: Mus musculus, Acomys russatus, Heterocephalus glaber, Xenopus laevis, Danio rerio, and Ambystoma mexicanum. (E)Protein docking models for predicted interactions between human SERCA2 and TRIM59-TM.

Immunoprecipitation (IP) and co-immunoprecipitation (co-IP) followed by liquid chromatography and mass spectrometry (LC-MS/MS) studies of SERCA2 from mammalian optic nerves corroborate this finding, showing that only NNAT and TRIM59 co-precipitate with SERCA2 in this tissue (Figure 4A). Additionally, the assays identified differences in SERCA2 and TRIM59 abundance between samples collected at developmental timepoints when the mouse optic nerve retains robust endogenous regenerative competence or undergoes regenerative failure after injury (Figure 4B), revealing a divergent regulatory framework for SERCA2 in these disparate regenerative paradigms (Figure 4C). Given that TRIM59 is intricately implicated in immune regulation and inflammatory signaling [69–71], these evolutionary modifications could suggest that ER Ca²⁺-dependent thermogenesis may have been co-opted to fine-tune immune and inflammatory competence in endotherms. However, the same Ca²⁺-driven inflammatory modulation, while evolutionarily beneficial, could impose a trade-off by promoting gliotic and growth suppressive responses that constrain regenerative scope, thereby advancing a thermoregulatory theory in which the evolution of endothermy mechanistically intersects with, and potentially hinders, tissue regeneration.

**Figure 4:**
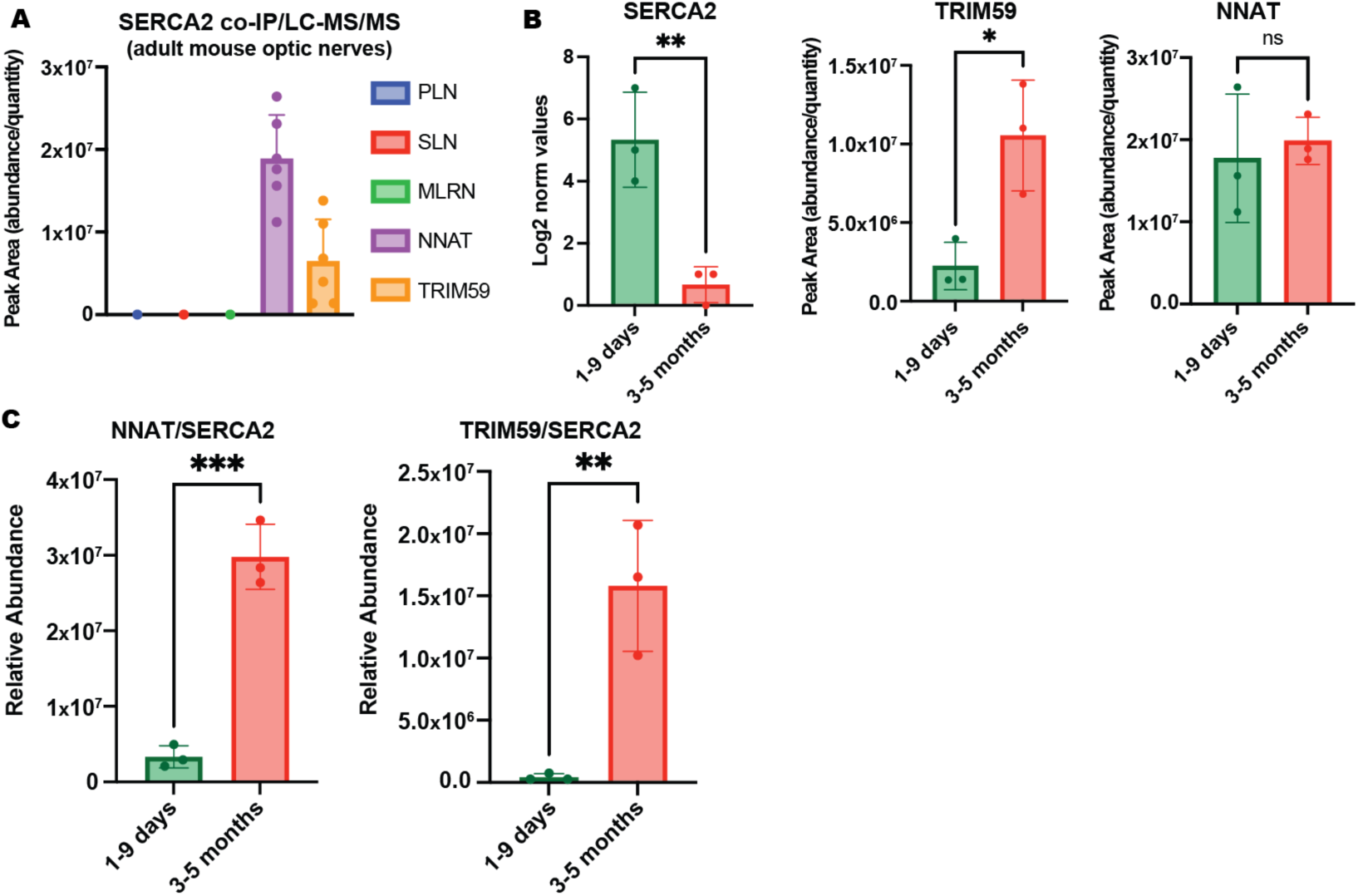
SERCA2 binding partners in the mammalian CNS. (A) SERCA2 co-IP and LC-MS/MS quantification of canonical SERCA2 regulators and TRIM59 in adult mouse optic nerves. (B) Mass Spectrometry quantification of SERCA2, TRIM59, and NNAT from mouse optic nerve tissue at two different developmental times. (C) Relative abundance of NNAT or TRIM59 to SERCA2 in mice optic nerves during the first 9 postnatal days, or at 3-5 months of age.

The central nervous system represents a convergence of several properties predicted by this framework to limit regenerative competence. Neurons are highly specialized, post-mitotic cells embedded within tightly regulated Ca²⁺ signaling architectures that couple electrical excitability to neurotransmitter release and metabolic demand. Unlike tissues capable of progenitor-driven replacement, functional restoration of neuronal circuits would require injured neurons to survive, reprogram aspects of terminal differentiation, and reestablish complex synaptic connectivity. Sustained intracellular Ca²⁺ elevations, particularly those linked to large and rapid ER Ca²⁺ release and impaired clearance, can activate transcriptional programs that favor inflammation, astrocytic reactivity, and extracellular matrix deposition rather than structural rebuilding of neuronal networks. Consequently, tissues with tightly orchestrated excitation–secretion or excitation–conduction systems may represent the contexts in which thermogenic Ca²⁺ dynamics architectures most strongly bias injury responses toward regenerative failure.

## Discussion

Here we advance a *Thermoregulatory Theory of Regenerative Scope*, proposing that the evolutionary acquisition and elaboration of composite thermogenic mechanisms imposed energetic and molecular constraints that progressively limited regenerative competence in highly metabolic tissues, with a particular focus on the adult mammalian central nervous system (CNS). Our framework integrates comparative physiology, developmental timing, and calcium-dependent signaling pathways to suggest that thermogenic specialization and regeneration are not merely coincidental traits, but mechanistically and evolutionarily intertwined.

Importantly, the framework proposed here does not mean to imply that endothermy constitutes a generalized systemic constraint on regenerative competence. Rather, we suggest that the evolutionary emergence of endothermic physiology created selective pressure for specialized molecular architectures that support rapid thermogenic responsiveness involving rapid calcium-dependent signaling networks whose regulatory consequences bias injury responses toward inflammation, fibrosis, or gliosis rather than functional regeneration, particularly in highly excitable tissues. Under this interpretation, endothermy provides the evolutionary context, whereas the mechanistic determinants of regenerative competence are local, tissue-specific properties of calcium-handling systems and injury-response programs. This distinction helps reconcile why regenerative capacity varies substantially among tissues within the same organism despite similar thermogenic output.

A central tenet of this theory is that the maturation of endothermic capacity, specifically mechanisms involving futile calcium cycling across the sarco/endoplasmic reticulum (SR/ER), directly interferes with the cellular processes required for functional regeneration. In mammals, shivering thermogenesis depends, in part, on uncoupling ATP hydrolysis from effective Ca²⁺ reuptake through SERCA via negative regulators such as sarcolipin and phospholamban. We hypothesize that removing or attenuating negative regulation of SERCA in non-regenerating tissues would re-couple ATP hydrolysis to rapid Ca²⁺ clearance, reduce cytosolic Ca²⁺ permanence after injury, and thereby enhance regenerative capacity. This prediction is consistent with experimental paradigms demonstrating that accelerated Ca²⁺ sequestration and restriction of ER Ca²⁺ release promote axonal regeneration, whereas prolonged intracellular Ca²⁺ elevation amplifies inflammatory transcriptional programs and fibrotic or gliotic remodeling [52, 55, 57].

Tissue-specific differences further support this mechanistic axis. Hepatocytes retain robust regenerative capacity despite their high metabolic demands. Notably, the liver lacks functional ryanodine receptors (RyR) [64, 65], potentially restricting rapid ER Ca²⁺ release and buffering against sustained Ca²⁺-driven inflammatory amplification. In contrast, cardiomyocytes and CNS neurons — cells that exhibit profound regenerative failure in adult mammals — are embedded in tightly regulated Ca²⁺ cycling architectures integral to excitation-contraction coupling, action potential propagation and thermogenic output. Hirose and colleagues [36] previously associated the postnatal acquisition of endothermic physiology in cardiomyocytes with loss of regenerative competence, linking thyroid hormone-dependent transitions to polyploidization and cell cycle exit. Thyroid hormones are established regulators of Ca²⁺ cycling proteins and ion transport complexes [72–74], including uncoupled ATPase activity by SERCA [75] providing a plausible endocrine bridge between systemic thermogenic maturation and intracellular calcium homeostasis. Another thyroid-specific calcium modulating hormone — calcitonin — is known to be highest during the same perinatal transition stages and decreases in serum concentration with age [76, 77]. Calcitonin has been shown essential for peripheral nerve regeneration[78, 79]. Extending this logic to the CNS, our data identifying NNAT and TRIM59 as candidate SERCA-associated interactors in mammalian optic nerve suggest that neural thermogenic specialization may likewise alter ER Ca²⁺ handling in ways that bias injury responses toward inflammation and gliosis rather than regeneration.

From an evolutionary perspective, the loss of regenerative capacity appears maladaptive. However, the molecular mechanistic burden of endothermy suggests that the trade-off may have conferred alternative survival advantages. Enhanced immune and inflammatory competence offers a compelling candidate. Sustained and rapidly mobilized Ca²⁺ flux is central not only to thermogenesis but also to activation of NF-κB–dependent transcription, inflammasome assembly, cytokine maturation, and reactive gliosis. The same ER Ca²⁺ dynamics that permit rapid thermogenic adaptation may therefore potentiate host defense in injury-prone, high-activity endotherms. In this view, regenerative suppression is not a passive loss but a systems-level consequence of prioritizing inflammatory efficiency and metabolic responsiveness. The co-option of ER Ca²⁺-dependent futile cycling to support immune surveillance and inflammatory amplification could have been selectively advantageous in endothermic lineages expanding to new and variable ecological niches.

While circumstantial at present, the convergence of evolutionary physiology, endocrine regulation, calcium signaling biology, and injury response supports a coherent and testable model. Distinguishing causation from evolutionary correlation represents a critical next step for evaluating this hypothesis. The theoretical framework generates several experimentally testable predictions. First, genetic or pharmacologic interventions that accelerate Ca²⁺ clearance or restrict ER Ca²⁺ release in adult mammalian CNS tissue should reduce injury-induced inflammatory amplification and enhance structural regeneration. Second, comparative studies across vertebrate species with divergent regenerative capacities should reveal systematic differences in SERCA regulatory architecture, RyR and/or IP_3_R expression patterns, and ER Ca²⁺ clearance kinetics. Third, developmental transitions associated with the maturation of thermogenic physiology, such as the postnatal acquisition of endothermy in altricial mammals, should coincide with measurable shifts in calcium-handling proteins, their regulatory architecture, and inflammatory signaling programs preceding the decline in regenerative competence.

By reframing regenerative failure not as an isolated deficiency but as a thermoregulatory byproduct of metabolic evolution, the *Thermoregulatory Theory of Regenerative Scope* provides a unifying lens through which disparate observations — ranging from neonatal mammalian plasticity to naked mole rat resilience and ectothermic regeneration — can be interpreted. Further mechanistic dissection of calcium-dependent thermogenic pathways in the CNS and other high-energy tissues will determine whether endothermy’s evolutionary success indeed came at the cost of regenerative potential.

